# Imaging biological tissue at extended depth using swept-source visible-light OCT

**DOI:** 10.64898/2026.10.16.759740

**Authors:** Weijia Fan, Roman Kuranov, Menglin Shi, Fengyuanshan Xu, Junfang Zhang, Hao F. Zhang

## Abstract

We present a swept-source visible-light optical coherence tomography (SS-vis-OCT) system with twice the imaging depth of prior SS-vis-OCT and improved sensitivity for imaging biological samples, achieving shot-noise-limited operation. Because second-harmonic generation is limited by the booster optical amplifier’s output power, the visible-light SS power is capped at 450 µW. Increasing it further would require a more powerful, but costlier, amplifier. Instead, we reduce the receiver noise floor by selecting a balanced detector with NEP approaching the thermal (Johnson–Nyquist) limit. Real-time numerical k-linearization increases the effective number of linear-in-wavenumber samples per sweep, further improving SNR and extending the imaging range. Together, these improvements yield a sensitivity of 89 dB and an imaging depth of 9 mm in air or 6.5 mm in tissue, enabling full anterior-segment imaging of *ex vivo* porcine eyes.

## 1. Introduction

Optical coherence tomography (OCT) provides non-invasive, depth-resolved, micrometer-scale imaging of biological tissue and has become a clinical standard in ophthalmology. Anterior segment OCT (AS-OCT) is routinely used to assess structures including the cornea, aqueous outflow system, anterior chamber, and ocular surface to manage corneal diseases, glaucoma, and other conditions [1-3].

Ultrahigh-resolution OCT extends the technique to resolving much finer structural detail either by shifting the source to a shorter center wavelength or by using a broader bandwidth. Ultrahigh-resolution OCT centered at 800 nm with a 170 nm bandwidth achieves an axial resolution of 1.2 µm in tissue. In human subjects, it has been shown to visualize and precisely measure corneal epithelial, Bowman’s layer, and paracentral stromal thickness [4, 5]. Visible-light OCT (vis-OCT), by contrast, uses a 510nm –610 nm wavelength range and provides comparable axial resolution, better layer contrast, and additional spectroscopic information in a regime where hemoglobin and melanin exhibit strong, well-separated absorption features. In small animals, visible OCT has resolved Schlemm’s canal and the limbal vascular network [6, 7].

All prior high-resolution anterior segment OCT, including vis-OCT, relies on spectrometer-based spectral-domain detection (SD-OCT). In this approach, the spatially dispersed light spectrum is convolved with the stepwise function of the spectrometer’s detection-array geometry, determined by pixel size, producing rapid depth-dependent sensitivity roll-off. The theoretical minimum roll-off, set by the Fourier transform of the finite pixel width, is about 1.44 dB/mm, independent of the central wavelength (see Appendix A). In practice, however, roll-off values below ∼5 dB/mm are rarely achieved because aberrations or imperfect optical alignment across the detector array limit performance of the spectrometers [8]. The wavenumber step between adjacent samplings sets the imaging depth of spectrometer-based OCT (Δ*k*_*lin*_) after linearization in wavenumber (*z*_*max*_ = *π*Δ*k*_*lin*_/2) , and for ultrahigh-resolution systems it reaches approximately 1.5 mm or less in tissue. In anterior segment imaging, because of corneal curvature, human ultrahigh-resolution OCT studies have been confined to the central cornea [3, 9, 10], not exceeding ±2-2.5 mm from the corneal apex in the lateral direction. However, wide-field imaging of the whole cornea and its transition to the sclera, as well as whole-anterior-chamber imaging, requires an imaging depth of 5.2 mm in tissue (7.2 mm in air), given an anterior chamber depth of 2.6–4.6 mm and a cornea thickness of 500–600 µm.

Swept-source OCT (SS-OCT) removes both constraints. Rather than dispersing the spectrum in space across a fixed pixel array, it encodes the spectrum in time by sweeping a narrowband laser and digitizing the signal on a single or balanced detector. Without the detector’s stepwise sampling, and with the light focused onto a single detector rather than imaged across an array, high-performance spectrometer optics are no longer required, and the sensitivity roll-off is instead governed mainly by the swept source’s instantaneous coherence length. This coherence length ranges from a few millimeters in FDML lasers [11] to meters in VCSEL sources [12, 13]. Widely used MEMS-based swept sources offer instantaneous coherence lengths of 10 mm–20 mm [14]. The maximum imaging depth depends on both the coherence length and the detection chain, and high-speed digitizers now acquire more than 6,000 samples per sweep, sampling the full tuning range at a fine wavenumber step. Together, these dramatically increase the imaging depth relative to spectrometer-based SD-OCT.

Because of NIR SS-OCT’s roll-off and imaging depth advantages, it has become the standard for whole-anterior-segment imaging, with systems capable of capturing the full cornea and anterior chamber in a single acquisition [15-17]. Developing swept-source vis-OCT (SS-vis-OCT) therefore promises the resolution and spectroscopic contrast of vis-OCT together with the extended imaging depth required for whole-anterior-segment imaging.

The previously reported SS-vis-OCT system, however, achieved only a limited sensitivity of 72 dB, insufficient for biological tissue imaging [18], and an imaging depth of 4.5 mm in air (3.3 mm in cornea), insufficient for whole anterior chamber imaging. Visible-light swept sources deliver limited output power, and the fraction available to the reference arm is insufficient to reach the shot-noise limit, even with the relative intensity noise rejection provided by balanced detection. Detector selection therefore becomes decisive in reaching the shot-noise limit under such power constraints [18]. However, commercial balanced receivers are overwhelmingly optimized for telecommunications bands, and visible-range silicon balanced detectors that combine high transimpedance gain, low input-referred noise, and bandwidths of hundreds of MHz adequate for 100 kHz-rate sweeps remain scarce.

Here, we show that detector noise, rather than source power, limits SS-vis-OCT sensitivity, and that an ultra-low-noise balanced receiver can overcome it. We present an SS-vis-OCT system that achieves shot-noise-limited operation at approximately 89 dB sensitivity, sufficient for biological-tissue imaging. This improvement comes from 1) selecting a balanced detector with noise-equivalent power approaching its thermal limit and 2) numerical k-linearization, which increases the effective number of samples per sweep. The same numerical k-linearization also decouples sampling density from the hardware-defined k-clock frequency, extending the imaging range from 4.5 mm to 9 mm in air, corresponding to 6.5 mm in tissue and exceeding the depth required for whole-anterior-chamber imaging. Together, these advances enable full anterior-segment imaging of an *ex vivo* porcine eye at depths beyond spectral-domain vis-OCT, setting the stage for clinical whole-anterior-segment vis-OCT.

## 2. Noise Analysis

When detecting photons, a photodetector operates as a current source *I*_*pd*_ = *RP*, where *R* (A/W) is the responsivity and *P* (W) is the incident optical power onto the photodetector. The transimpedance amplifier converts this current to a voltage *V* = *Z*_*T*_*I*_*pd*_ to be digitized by the analog-to-digital converter (ADC), where *Z*_*T*_ (V/A) is an effective transimpedance gain.

Hereafter, noise is expressed as a voltage variance (*V*^2^, which at the ADC output comprises two optical and three electronic terms:

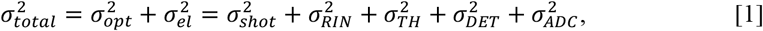

where the shot noise, RIN noise, and thermal noise voltage variances are [19-21]:

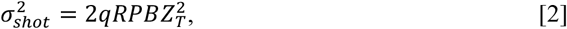

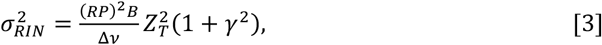

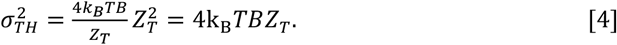

Here, *P* = *P*_*ref*_ + *P*_*samp*_ , where *P*_*ref*_ and *P*_*samp*_ are the respective power values returned from the reference and sample arms, incident on each photodiode of the balanced detector. Since *P*_*ref*_ ≫ *P*_*samp*_ in biological tissue imaging, this reduces to *P* ≈ *P*_*ref*_ , which is adopted throughout this work. In the above equations, *q* (C) is the electron charge, *B* (Hz) is the balanced detector bandwidth, Δ*ν* (Hz) is the optical linewidth, *γ* (unitless) is the excess noise factor, *k*_*B*_ (J/K) is Boltzmann’s constant, and *T* (K) is temperature. The electronic noise term collects all light-power-independent contributions of the detection chain: the Johnson–Nyquist thermal noise of the transimpedance stage, 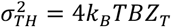, plus the detector amplifier noise 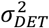 and the ADC electronic noise 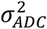.

The OCT interferogram signal is given as 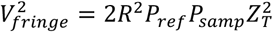, hence, the system SNR after Fourier transformation is

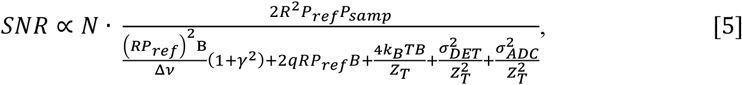

where *N* is the number of samples digitized per sweep. During the discrete Fourier transformation, the interference signal at a given depth is coherently summed, amplifying the signal power by *N*^2^, while the noise adds incoherently, scaling the noise power by *N*. The SNR is therefore proportional to *N*.

To approach the shot-noise limit, the RIN and receiver noise terms must be minimized relative to the shot noise. The RIN scales with 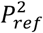 and thus grows faster with increasing reference power than the shot noise, dominating at high reference-arm powers. Balanced detection suppresses RIN noise, but doubles both the shot and receiver noise contributions because the two detector outputs add independently.

With the removal of RIN, shot noise limited condition is achieved when

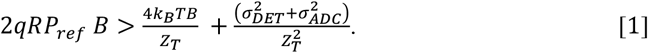

This means, under limited *P*_*ref*_ , the electronic noise must be reduced. There are two considerations. First, the electronic noise 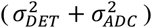 is bounded below by 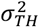, the fundamental physical floor of the detection chain. A detector whose amplifier noise 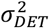 is negligible against 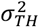 operates at this limit and is therefore preferred. Second, the transimpedance gain *Z*_*T*_ sets the weight of the receiver noise in the total noise floor. Increasing *Z*_*T*_ amplifies the signal and the shot noise as 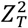, whereas the thermal noise increases only as *Z*_*T*_. Referred to the input optical power, a detector with a higher *Z*_*T*_ , therefore, suppresses the receiver noise’s relative contribution and lowers the reference-arm power required to reach the shot-noise-limited regime. However, *Z*_*T*_ cannot be increased indefinitely, as higher transimpedance gain narrows the detection bandwidth, which must remain sufficient to capture the full fringe frequency bandwidth. In practice, the detector’s noise-equivalent power captures these requirements and forms the basis for detector selection in Section 3.2.

## 3. Method

### 3.1 Experimental setup

The SS-vis-OCT used a visible-light swept source (SS) based on second harmonic generation using a periodically poled lithium niobate (PPLN) crystal [22]. Briefly, a near-infrared (NIR) SS (central wavelength 1310 nm, bandwidth 100 nm, sweeping rate 100 kHz) was amplified by a booster optical amplifier (BOA-1280-50PM-500, Innolume, Germany; center wavelength 1280 nm, bandwidth 50 nm), then frequency-doubled through the PPLN to produce a visible SS centered at 640 nm with 25 nm bandwidth. The BOA saturates at 300 mW, yielding 450 μW of SHG power.

The output of the second-harmonic-generated visible light SS was sent into an interferometer, as shown in Fig. 1. The interferometer followed a fiber-based Mach-Zehnder configuration for balanced detection. A 50/50 fiber coupler (FC1) split the source light into sample and reference arms. In the sample arm, a collimating lens (CL, focal length = 10 mm) collimated the light and directed it onto a scanning galvanometer mirror pair (GM) before an objective lens (focal length = 50 mm) focused it onto the sample. Optical power at the sample was 220 μW, and light reflected from the sample was collected back through the same path in the sample arm. In the reference arm, light passed through a polarization controller (PC2) and traversed a free-space path formed by two retroreflecting mirrors (RM) mounted on a translation stage, allowing adjustment of the reference arm path length to match that of the sample arm. Light returning from the sample and reference arms was recombined at 50/50 fiber coupler (FC2) and directed to a balanced photodetector.

**Figure 1.**
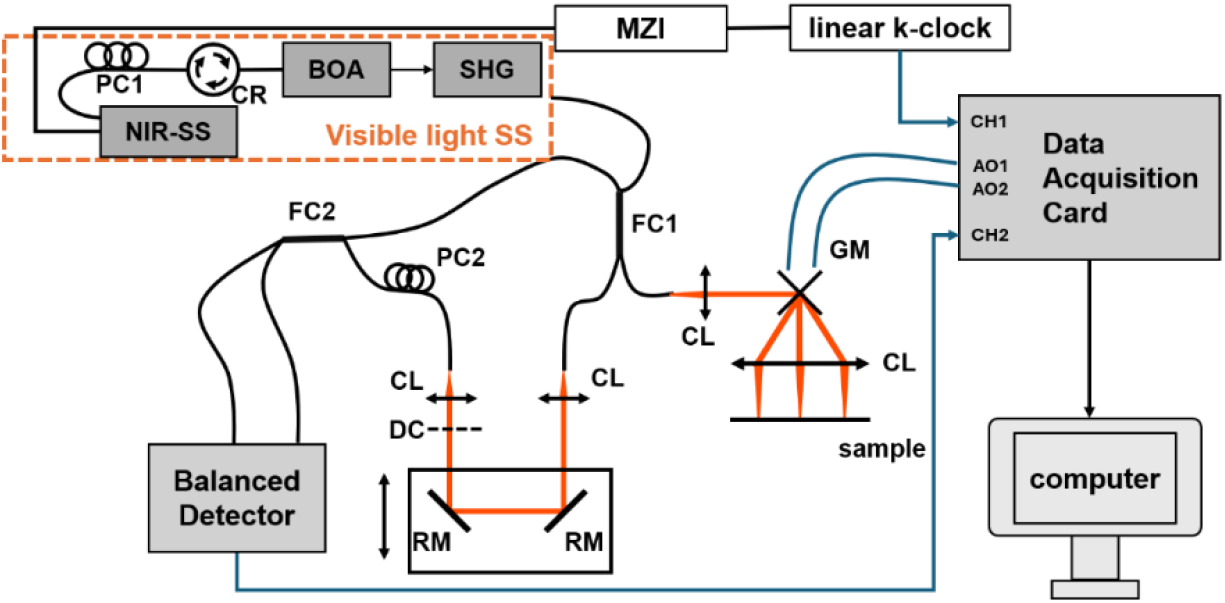
Schematic of the SS-vis-OCT experimental system. AO: analog output; BOA: booster optical amplifier; CH1, CH2: ADC channels of the DAQ; CL: collimating lens; CR: circulator; DC: dispersion compensation glass; FC: fiber coupler; GM: galvo mirrors; MZI: Mach-Zehnder interferometer; PC: polarization controller; RM: reflecting mirrors; SHG: second harmonic generator.

We used a data acquisition (DAQ/ADC) card (SA220P, Acqiris, Switzerland) to control the galvanometer scan and digitize the balanced-detector signal. The sweep trigger output from the NIR SS was directed to the external trigger channel of the SA220P, and the galvanometer scan pattern was driven directly by two analog output channels (AO1, AO2) of the same card. Data acquisition was likewise initiated on each trigger, so that beam scanning and signal digitization were generated and triggered within a single device, inherently synchronizing the two.

A fraction of the NIR SS output was tapped to an internal Mach–Zehnder interferometer, generating an auxiliary k-clock whose frequency is proportional to the instantaneous sweep rate and optical path delay between the Mach-Zehnder arms. Rather than using this k-clock to trigger the digitizer directly, we digitized the k-clock and the balanced-detector fringe simultaneously on two DAQ input channels at 2 GS/s with 14-bit depth, sharing a common time base. Acquisition was gated to the 3.2 μs forward portion of each 10 μs sweep spanning the spectral range used for visible SHG, yielding approximately 6528 samples per sweep. At the k-clock’s maximum specified frequency of 317 MHz, the digitization yields roughly 6 samples per k-clock cycle. Resampling indices for uniform wavenumber spacing were then computed numerically from the unwrapped Hilbert-transform phase of the digitized k-clock and applied to the interferogram. The card’s on-board FPGA performed this operation in real time during acquisition [23]. The k-linearized interferograms were streamed continuously to the host computer and buffered in on-board DDR memory, eliminating post-acquisition resampling. Dispersion compensation was then performed using the sub-band registration method to determine optimal second- and third-order coefficients [24], followed by FFT to retrieve A-line information and reconstruct OCT images.

### 3.2 Balanced detector selection

In practice, a balanced detector’s receiver noise floor is specified by its noise-equivalent power 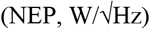. The thermal noise of the transimpedance sets its lower bound as

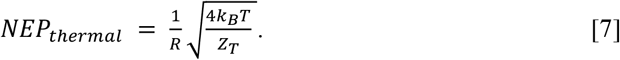

A lower absolute NEP lets shot noise exceed the receiver noise at a lower reference-arm power, and a detector reaches this thermal bound only when its NEP approaches *NEP*_*thermal*_ . However, NEP is not only a function of the detector’s efficiency but also a function of the detector’s gain, and a detector with higher efficiency and larger gain can have larger NEP.

We evaluated each detector by two quantities. The first is the absolute NEP: the detector with the lowest absolute NEP reaches shot-noise-limited operation at the lowest reference-arm power. The second is the detector’s gain-independent dimensionless ratio

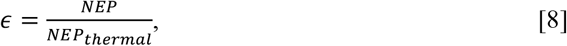

which indicates the portion of the detector’s NEP attributed to fundamental thermal noise. Values of *ε* approaching unity indicate a thermal-noise-limited detector, and larger values indicate excess electronic noise.

We compared two balanced detectors with comparable bandwidth: a Thorlabs PDB435A (Thorlabs, NJ, USA) and an LD-PD UBD-400M-B (LD-PD PTE LTD, Singapore). Table 1 summarizes the relevant specifications: detector material, wavelength range, bandwidth, transimpedance gain, and specified NEP from the manufacturer datasheets, and responsivity read from the provided curves at 640 nm. The thermal-limited NEP and *ε* were computed from the datasheet transimpedance gain and responsivity via Eq. 8. The higher transimpedance gain of the UBD-400M-B yields a lower thermal-limited NEP, and its lower *ε* indicates that this floor is nearly achieved. We therefore selected the UBD-400M-B, enabling shot-noise-limited operation within the available source power.

**Table 1.** Specifications of two candidate balanced detectors.

| Parameter | PDB435A | UBD-400M-B |
| --- | --- | --- |
| Detector material* | Si | Si |
| Wavelength range* (nm) | 320–1000 | 400–1100 |
| Bandwidth* $B$ (MHz) | 350 | 400 |
| Responsivity* $R$ (A/W) @640nm | ~0.4 | ~0.35 |
| Transimpedance gain* $Z_T$ (V/A) | $1.0 \times 10^4$ | $7.2 \times 10^4$ |
| $NEP^*$ (pW/$\sqrt{\text{Hz}}$) | 32.3 | 5 |
| $NEP_{thermal}^{**}$ (pW/$\sqrt{\text{Hz}}$) | 4.55 | 1.37 |
| $\epsilon = NEP/NEP_{thermal}$ | 7.1 | 3.7 |

\* - parameter is from the detector's spec sheet
\*\* - parameter is calculated using Eq. 8

### 3.3 System sensitivity measurement

We measured the noise floor and system SNR as a function of reference-arm power for both balanced detectors to confirm shot-noise-limited operation. For the noise floor measurement, the sample arm was blocked, and a variable attenuator was placed in the reference arm to control *P*_*ref*_ . At each power level, we acquired the reference arm spectrum, removed the direct current (DC) component, and recorded the standard deviation of the resulting spectrum centered at 0 V as the noise. With both arms blocked, we measured the system’s total electronic noise independently, including the thermal noise of both detectors and the ADC card. To measure the ADC noise, we ran the digitization with the detector powered off.

For the sensitivity measurement, a neutral density filter of known attenuation *D* was placed in the sample arm, and a variable attenuator in the reference arm was adjusted to maximize *P*_*ref*_ without saturating the fringes [25]. The SNR was calculated as

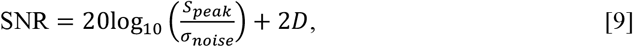

where *S*_*peak*_ is the A-line PSF peak amplitude, and *σ*_*noise*_ is the standard deviation of the FFT of the mean-subtracted reference arm spectrum.

### 3.4 Imaging depth validation

We measured the system sensitivity roll-off using a mirror placed at the sample-arm focal plane and attenuated by a neutral-density filter. We set the reference-arm delay in known increments to position the resulting PSF peak at a series of optical path-length differences spanning the full depth range. At each position, the k-linearized and dispersion-compensated interferogram was Fourier-transformed, and the peak amplitude and axial width of the PSF were extracted. We calculated the roll-off at 1 mm as the drop in peak amplitude (in dB) between the PSF at 1 mm and that near zero delay.

To further verify the imaging depth, we imaged a pyramid phantom of known height [22]. We 3D-printed the phantom in polymer as a stepped pyramid with 600 μm step heights on one side and 200 μm step heights on the other, with overall dimensions of 12 mm (L) × 12 mm (W) × 12 mm (H). We imaged it over a 12.5 mm lateral field of view with 1024 A-lines per B-scan.

### 3.5 Anterior segment imaging and sample preparation

For anterior segment imaging, we used a porcine eye because its globe size and anterior segment geometry are comparable to those of the human eye [26, 27]. We obtained the porcine eyes from a USDA-inspected local butcher shop and transported them to the laboratory on ice. Upon arrival, we stored the eyes at 4°C until imaging. We imaged the eyeballs within 72 hours after harvesting. Before imaging, we visually inspected the globe to confirm the eyeball was intact and the cornea remained visibly clear. During imaging, we also placed a commercially available soft contact lens (-3.5 diopters, 8.8 base curve; ACUVUE OASYS, Johnson & Johnson Vision, Jacksonville, FL) on the corneal surface as part of the imaged sample. The contact lens was not used for optical correction or hydration control.

SS-vis-OCT images were acquired over a 20 mm lateral field of view, with each B-scan consisting of 2048 A-lines. This scan configuration assessed the SS-vis-OCT’s ability to capture the curved anterior segment surface over an extended imaging depth. For comparison, we also imaged the same sample using a home-built SD-vis-OCT system described previously [28]. The SD-vis-OCT system used a supercontinuum laser source filtered to 510–610 nm and a spectrometer with 2048 pixels (OpticentHealth, Evanston, IL). The SD-vis-OCT system had an imaging depth of 1.5 mm in air. Because of this limited imaging depth and the curved geometry of the porcine eye, we used a smaller lateral field of view to highlight SD-vis-OCT limitations and for comparison imaging. Therefore, SD-vis-OCT B-scans were acquired over a 10 mm lateral range with 2048 A-lines per B-scan.

## 4. Results

### 4.1 Noise and sensitivity

Figure 2 shows the results for both balanced detectors, with blue curves representing the Thorlabs PDB435A and red curves representing the UBD-400M-B. In Fig. 2a, the solid lines show the theoretical total noise, the horizontal dashed lines show the constant electronic noise floor of each detector and ADC card, and the curved dashed lines show the theoretical shot noise contribution, which scales as 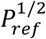. The measured data points (circles) match the theoretical predictions well. The PDB435A exhibits a significantly higher thermal noise floor, such that the measured noise remains thermally dominated across the entire practical range of reference arm powers, never reaching the shot noise limit. In contrast, the UBD-400M-B has a substantially lower thermal noise floor, and the measured noise transitions to the shot-noise-limited regime at moderate reference-arm powers accessible in our system.

**Figure 2.**
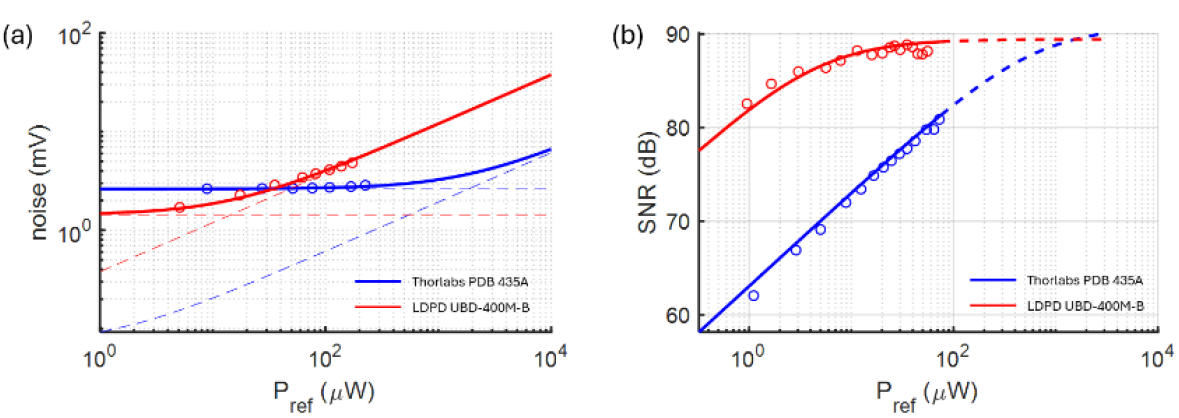
(a) Calculated (lines) and measured (circles) noise using balanced detectors PDB 435A (blue) and UBD-400M-B (red) at different *P*_*ref*_. The horizontal dashed lines mark the electronic noise for the two detectors, respectively, and the curved dashed lines show the theoretical shot noise contribution. (b) Calculated (lines) and measured (circles) SNR using balanced detector PDB 435A (blue) and UBD-400M-B (red) at different *P*_*ref*_ levels. The solid line covers the maximum achievable power in the current system, and the dashed line extends the prediction using the theoretical model.

In Fig. 2b, the solid lines show the theoretical SNR up to the maximum reference-arm power reaching the detector in our system. In contrast, the dashed lines extend the prediction to higher powers. In the shot-noise-limited regime, SNR saturates as reference power increases; in the presence of residual RIN, it instead rolls off at high powers. The measured data points align well with theoretical predictions. The UBD-400M-B reaches a constant SNR of 89 dB starting at *P*_*ref*_ = 20 µW, confirming shot-noise-limited operation within our available power range. The absence of a high-power SNR roll-off is consistent with effective common-mode RIN suppression by the balanced detector over this range. In contrast, the PDB435A’s SNR continues to rise across the entire available reference arm power range, indicating that light-power-independent electronic noise 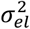 still dominates and shot-noise-limited operation has not been reached. Extrapolating from the simulation, the PDB435A would require a *P*_*ref*_ of ∼3000 µW to reach the shot-noise limit, corresponding to a source visible light SS power of over 12 mW, assuming no coupling loss in the reference arm, far exceeding the BOA output ceiling. This confirms that detector noise, rather than optical power, limited the previous system, and that optimizing detector performance is a viable path to shot-noise-limited visible-light SS-OCT.

### 4.2 Imaging depth validation

Figure 3a shows the point-spread functions (PSFs) recorded at successive mirror depths across the imaging range. The peak signal fell by 0.39 dB over 1 mm and by 30 dB across the full 9 mm range. The 6 dB roll-off depth was 3.9 mm. Fig. 3b shows the PSF measured at 1 mm depth, with a full width at half maximum of 9.5 μm in air, giving the system axial resolution below 10 μm in air and 6.9 μm in cornea (n = 1.38).

**Figure 3.**
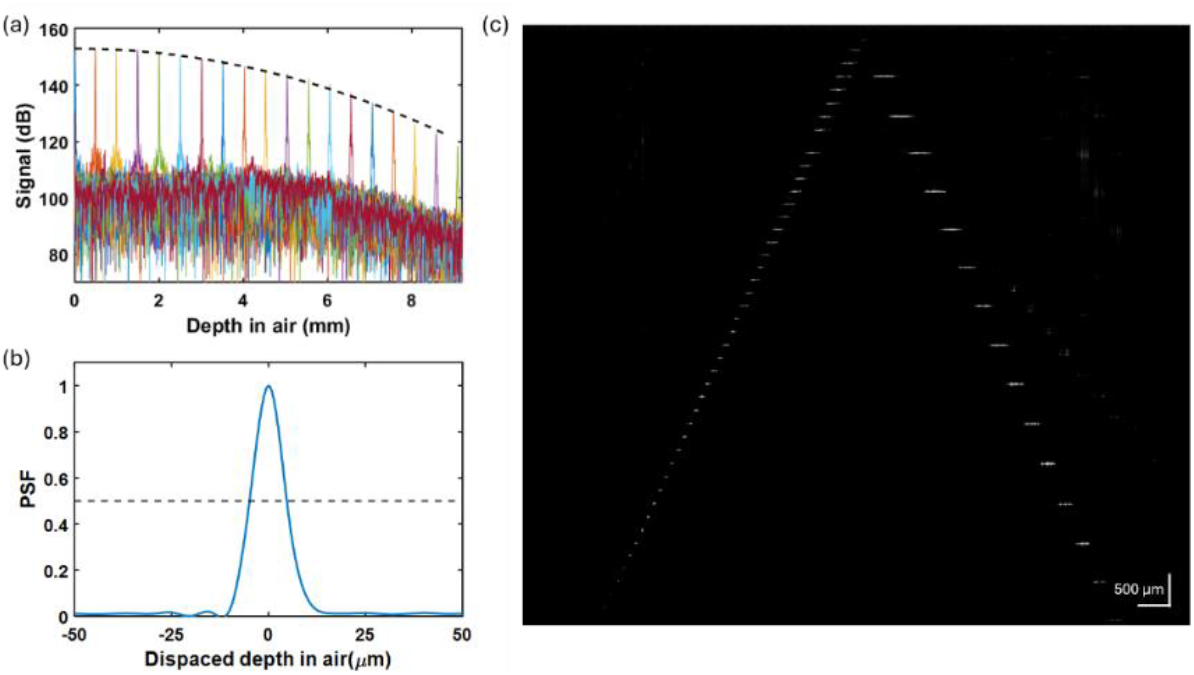
Imaging depth validation result. (a) The sensitivity roll-off of the SS-vis-OCT system. The dashed line corresponds to the roll-off fit. (b) PSF at 1 mm depth position with the measured full width at half-maximum (FWHM) being 9.5 µm. (c) Representative B-scan of a pyramid phantom.

Figure 3c shows a representative B-scan of the 3D-printed pyramid phantom. Fifteen consecutive steps of 600 µm height are resolved in the cross-section and remain distinguishable at the deepest steps despite the reduced signal there. This confirms the system’s imaging depth is over 9 mm.

### 4.3 Cornea imaging

Figure. 4 shows representative B-scans of an *ex vivo* porcine eye. The images compare the SS-vis-OCT and SD-vis-OCT systems under two sample conditions: without a contact lens and with a contact lens on the corneal surface.

**Figure 4.**
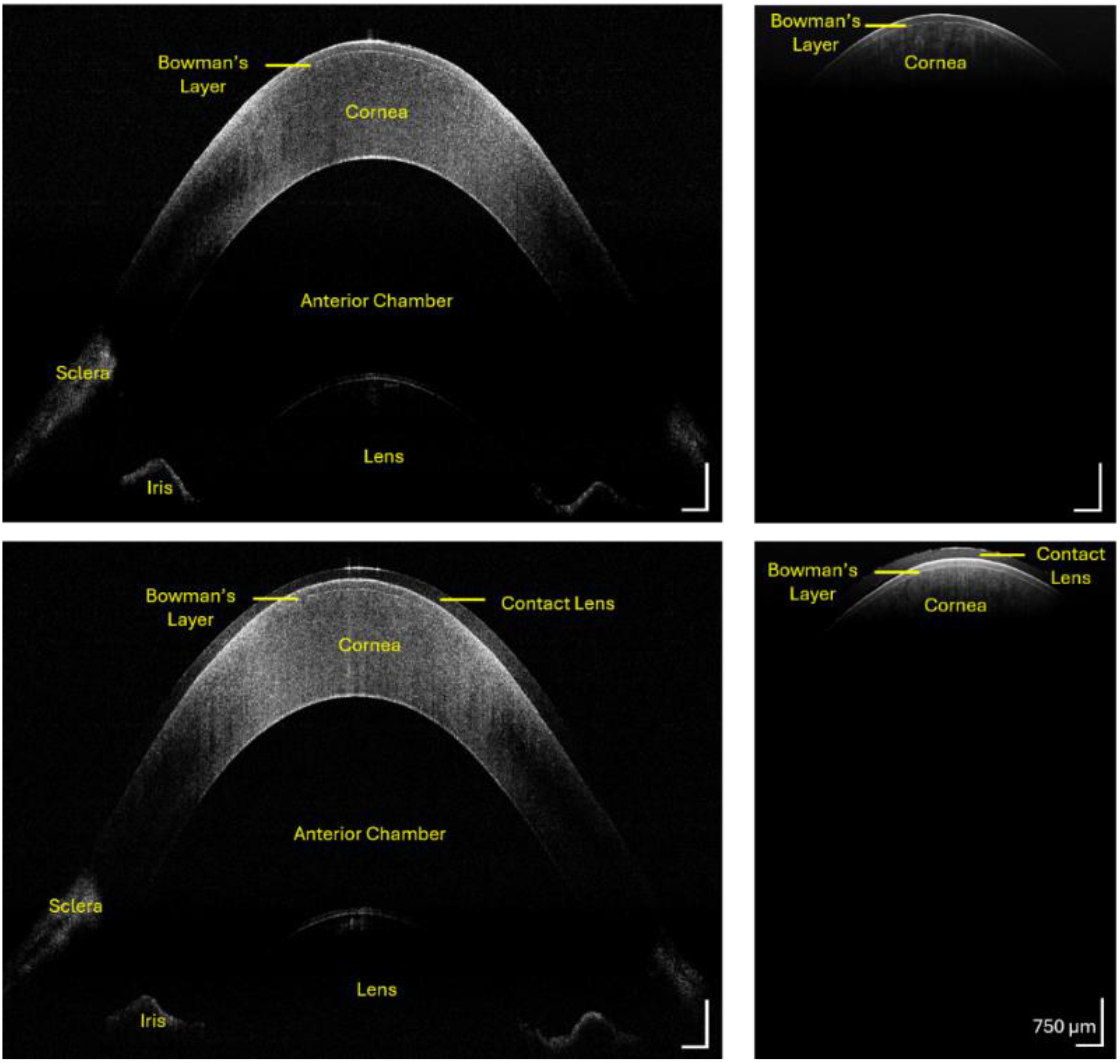
*Ex vivo* porcine anterior segment imaging with SS-vis-OCT and SD-vis-OCT. (a) SS-vis-OCT and (b) SD-vis-OCT B-scans without a contact lens. (c) SS-vis-OCT and (d) SD-vis-OCT B-scans with a contact lens. The SS-vis-OCT system provides an imaging depth of 9 mm in air, clearly resolving the cornea, iris, and anterior lens surface, while the SD-vis-OCT system is limited to 1.5 mm in air, capturing only the anterior corneal surface. Scale bar: 750 µm.

Without a contact lens, the SS-vis-OCT image (Fig. 4a) clearly resolves the front and back surfaces of the cornea, Bowman’s layer, the iris, and the anterior surface of the lens. The SD-vis-OCT system (Fig. 4b), limited to 1.5 mm in air (1.1 mm in tissue), captures only the anterior corneal surface and resolves Bowman’s layer at 1.3 μm axial resolution but cannot reach the deeper structures. With a contact lens, the SS-vis-OCT image (Fig. 4c) clearly resolves both surfaces of the contact lens and the full cornea beneath it. In contrast, the SD-vis-OCT image (Fig. 4d) shows only the central portion of the contact lens anterior surface; as the contact lens curves away from the zero delay, deeper portions exceed the 1.1 mm imaging depth and are lost. SS-vis-OCT thus preserves layer-level corneal detail from high-resolution vis-OCT while extending coverage to the wide-field, full-depth anterior segment in a single B-scan, a combination SD-vis-OCT cannot achieve. Resolving both the full contact-lens geometry and its position relative to the corneal surface suggests potential applications of SS-vis-OCT in contact-lens inspection and in evaluating on-eye lens fit.

## 5. Discussion

### 5.1 K-linearization method for extending imaging depth

As shown in Section 3.2, measured sensitivity over depth shows a 6 dB roll-off over an imaging range of 3.9 mm, with usable signal extending beyond 9 mm, as confirmed by pyramid phantom imaging. This extended depth beyond the previous SS-vis-OCT system [22] results from performing real-time k-linearization numerically rather than externally clocking the digitizer from the k-clock. In hardware-triggered SS-OCT, acquisition is limited to one sample per physical clock transition, so the k-sampling density, and hence the maximum imaging depth, is bounded by the k-clock frequency. In the previous SS-vis-OCT implementation, even with hardware frequency doubling before the clock input, the imaging depth was limited to 4.5 mm, insufficient to capture the full anterior segment and short of what the source coherence length could support. Instead, deriving the resampling points in software decouples the k-sampling density from the physical clock rate and allows efficient sub-external-clock-rate resampling. This yields a 9 mm imaging depth, a two-fold improvement using the same source and k-clock. Sub-external-clock-rate resampling also increases the number of effective digitization points N per sweep, theoretically raising the SNR through its N-dependence (Eq. 5) and thus contributing to the measured sensitivity improvement alongside shot-noise-limited operation. Notably, the digitizer’s on-board FPGA performs this digital resampling in real time on every sweep, removing the computational burden from post-processing.

With the clock limitation removed, the detection bandwidth, the digitizer, and the source coherence length jointly set the maximum usable depth in SS-OCT. The hard limits of the detection are not reached here. The fringe frequency at 9 mm lies below the 400 MHz detector bandwidth, and the 2 GS/s digitizer satisfies the Nyquist requirement both for sampling the fringe signal and for sampling the k-clock signal. The observed sensitivity roll-off is consistent with the visible source’s finite coherence length. Residual errors in the software k-linearization may also contribute at the largest depths, since resampling fidelity depends on the sampling-rate-to-clock-frequency ratio and clock stability. Reducing this roll-off therefore calls for both a longer-coherence-length NIR seed and a higher digitizer sampling rate relative to the clock frequency. Commercial MEMS-VCSEL swept sources offer coherence lengths exceeding 100 mm [13, 29], which, combined with high-fidelity resampling, would potentially extend the visible coherence length and imaging range well beyond the present system.

The extended range is significant for the clinical prospects of vis-OCT. Prior vis-OCT has demonstrated unique spectroscopic contrast and fine structural detail in small-animal models [6, 7]. Still, SD-vis-OCT is typically limited to 1.1 mm of imaging depth in tissue, extending to roughly 1.7 mm in water with full-range methods [30], which is insufficient for many anterior-segment applications. The 9 mm in air (6.5 mm in tissue, n = 1.38) imaging range demonstrated here, obtained in a porcine eye of dimensions comparable to the human eye [26,27], indicates that vis-OCT can reach depths relevant to full-depth, wide-field anterior-segment imaging, supporting its potential for clinical translation.

### 5.2 Shot-noise-limited detection under source-power constraints

The central constraint of SHG-based visible SS-OCT is a dual power ceiling on both arms of the interferometer. We capped the reference-arm power because second-harmonic conversion saturates with the BOA output, limiting usable visible power to 450 µW. This differs from conventional NIR OCT, where swept sources deliver tens of milliwatts, reference power is abundant, and shot-noise-limited operation is readily reached. Ocular safety and aversion limits separately cap the sample-arm power. Together, these make shot-noise-limited operation essential, not incidental. First, with reference power capped, the regime is reached only if the receiver-to-shot-noise crossover falls at low reference power, achievable only with a sufficiently low-noise detector. Second, with sample power capped, sensitivity at the shot-noise limit is fixed by the sample-arm power and A-line rate. At ∼200 µW and 100 kHz, this ceiling must exceed the ∼85 dB required for biological-tissue imaging, leaving no margin for residual receiver noise. Therefore, to meet this sensitivity requirement, we focused on detector selection rather than increasing power. Using the same BOA-limited source as the previous system, we raised the sensitivity by 15 dB to 89 dB, confirming that receiver noise, not optical power, was the limiting factor.

This motivates a detector-customization criterion tailored to power-constrained operation. The detector that reaches shot-noise-limited operation at the lowest reference power is the one with the lowest absolute input-referred NEP. The ratio *ϵ* introduced here indicates how much of that NEP is unavoidable thermal noise versus excess electronic noise. The UBD-400M-B combines a low absolute NEP with *ϵ* ≈3.7, so it reaches the shot-noise limit at low power and sits near its physical floor, whereas the PDB435A (higher NEP, *ϵ* ≈ 7.1) is dominated by excess electronic noise and remains receiver-noise-limited across the accessible range.

Notably, among Thorlabs’ 400 MHz OCT detectors, the visible PDB435A (*ϵ* ≈7.1) sits about twice as far from its thermal limit as the NIR PDB470C and PDB471C (PDB470C, operating at 1300nm, *ϵ* ≈ 3.7; PDB471C-AC, operating at 1060nm, *ϵ* ≈ 3.2), indicating that the visible Thorlabs part carries more excess electronic noise than its NIR counterparts of equal bandwidth. The UBD-400M-B (*ϵ* ≈3.7) is competitive with the best NIR detectors. The scarcity of low-noise visible balanced detectors therefore reflects the application’s novelty rather than any technology limitation. Commercial fiber-coupled balanced photodetectors are predominantly built around InGaAs photodiodes for the near-infrared, a legacy of the telecommunications and NIR swept-source OCT markets that drove their development. In the visible range, no comparable demand existed. Prior low-noise vis-OCT was demonstrated almost exclusively in the spectral domain using dual calibrated spectrometers rather than discrete balanced detectors. As one of the first swept-source vis-OCT systems, this work operates where few low-noise commercial visible balanced detectors exist, yet the available detector still approaches its thermal-noise limit and reaches shot-noise-limited operation comparable to established NIR detectors. By establishing swept-source vis-OCT as a viable tissue-imaging modality, this work may help create the demand that drives development of dedicated ultra-low-noise visible balanced detectors.

### 5.3 Towards high power and higher resolution visible SS-OCT

Two future developments would substantially extend the capability of the present architecture, and both rest on components that already exist. The first is optical power at the sample. Shot-noise-limited detection ensures that no sensitivity is lost to the receiver, but it does not increase the number of photons returning from the sample; in strongly scattering tissue, the signal attenuates with depth until sample power becomes the penetration constraint. Where the sample arm is not safety-capped, as in anterior-segment imaging, penetration can be improved by raising the incident power. Where ocular laser safety limits incident power, as in retinal imaging, sensitivity must instead be improved through collection efficiency by replacing the current 50/50 fiber coupler (FC1) with a 90/10 or 95/5 split to direct more returning light to the detector at fixed tissue-incident power. Both routes require greater source power and because the system is shot-noise-limited, this added power converts directly into sensitivity rather than overcoming electronic noise. Broadband high-power booster optical amplifiers developed for telecommunications, with ∼100 nm bandwidth centered at 1310 nm and saturation powers approaching 1 W [31] would increase the available visible power by roughly an order of magnitude. Adopting such an amplifier in the visible light SS is therefore a clear path to the deeper penetration and higher sensitivity needed for retinal and deep-tissue SS-vis-OCT.

The second is the choice of operating wavelength. Because the visible sweep is generated by frequency doubling, its center wavelength and bandwidth are inherited directly from the NIR seed, and the present 640 nm output reflects the 1310 nm seed rather than any optimum for visible-light OCT. Seeding at 1050 nm would instead produce a visible sweep centered near 525 nm, aligning the source with the around 560 ± 50 nm range used in conventional spectral-domain vis-OCT. This shift improves axial resolution for a given fractional bandwidth, and it places the sweep across the hemoglobin absorption features that underpin the spectroscopic oximetry contrast distinctive to visible-light OCT. The 1050 nm window also benefits from a more mature amplifier ecosystem where semiconductor optical amplifiers at these wavelengths are commercially available with both broader gain bandwidth and higher saturation power. A swept-source implementation at these wavelengths would therefore combine vis-OCT’s spectroscopic contrast with the depth range and detection sensitivity demonstrated here.

## 6. Conclusion

We demonstrated a visible-light swept-source OCT system operating at the shot-noise limit by reducing the receiver noise floor rather than increasing source power. Because available power is limited by both the SHG conversion ceiling and ocular safety constraints specific to visible light, detector selection, rather than source engineering, becomes decisive for reaching the sensitivity required for biological-tissue imaging. Evaluating candidates by their input-referred NEP and the ratio of that NEP to its thermal limit, we selected a balanced detector whose receiver noise is much closer to the physical thermal floor than that of a conventional device with comparable bandwidth. This, together with increasing the number of digitization events, yielded shot-noise-limited operation within the available power budget and a ∼15 dB sensitivity improvement to 89 dB, confirming that electronic noise, not optical power, was the limiting factor. Additionally, performing k-linearization numerically rather than clocking the digitizer directly further decoupled the sampling density from the physical clock rate, extending the imaging range from 4.5 mm to 9 mm.

In conclusion, these improvements enabled visualization of the full anterior segment of an *ex vivo* porcine eye in a single B-scan, indicating potential for high-resolution, full-depth anterior-segment imaging at clinically relevant scales. More broadly, the detector-selection criterion developed here applies to any power-constrained interferometric detection problem, with the absolute NEP setting the reference power required for shot-noise-limited operation and its ratio to the thermal limit distinguishing unavoidable from excess noise. Seeding the same SHG architecture at a shorter pump wavelength would place the visible sweep within the range used in conventional spectral-domain vis-OCT, opening the possibility of combining its distinctive spectroscopic contrast with a depth range suited to whole-anterior-segment imaging.

