## Appendix A for "Imaging biological tissue at extended depth using swept-source visible-light OCT"

### Appendix A. Full-bandwidth theoretical minimum roll-off in FD-OCT

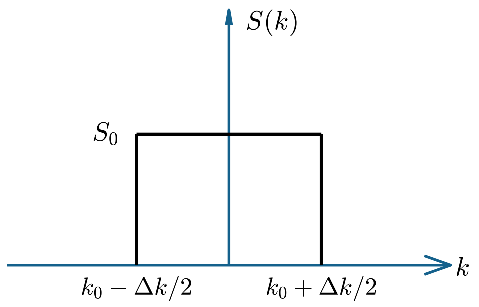


Figure1. Power spectrum at a single detector pixel. For simplicity, the power spectrum is constant over the pixel. We assume that the wavenumber bandwidth at this specific pixel is close to the effective wavenumber step over the pixels: $\Delta k = 2\pi/N\ (1/\lambda_{start}\ - 1/\lambda_{end}\ ) = 0.99\ mm^{-1}$ , where N=2048 is the number of pixels of the array detector; $\lambda_{start}$ =0.51 µm is the shortest wavelength of the spectrometer; and $\lambda_{end}$=0.61 µm is the longest wavelength of the spectrometer.

As shown by Yun et al [1] for full bandwidth and Rubinoff et al [1] for full and spectrally dependent bandwidths, assuming an ideal spectrometer in which roll-off from optical aberrations and finite spot size is negligible, the roll-off imposed by sampling of the spectrum onto the detector pixels is the Fourier transform of the power spectrum over the effective wavenumber band incident on a single pixel (Eq. 1). This constitutes the pixel sampling limit, the minimum roll-off attainable when the finite pixel width is the sole remaining contribution:

$$\frac{R(z)}{R(0)} = \left| \int_{k_0-\frac{\Delta k}{2}}^{k_0+\frac{\Delta k}{2}} S_0 e^{i2kz}\, dk \right| = \frac{sin(\Delta kz)}{\Delta kz} = sinc(\Delta kz), \qquad [1]$$

Hereafter, we refer to the roll-off described by equation 1 as theoretical roll-off. At the depth of 1 mm, the normalized theoretical roll-off for the present FD-OCT configuration is given by:

$$R(1\ mm)_{norm} = 20log10(\frac{R(z=1mm)}{R(0)}) = -1.44\ dB\ , \qquad [2]$$

The pixel sampling roll-off limit is therefore 1.44 dB/mm. For a specified axial resolution, this limit is independent of the central wavelength $\lambda_o$:

$$\frac{R(z)}{R(0)} = sinc(\Delta kz) = sinc\left(\frac{4ln2\cdot z}{N\cdot\delta z}\right), \qquad [3]$$

where $\delta z = \sigma\cdot\lambda_o^2/\Delta\lambda$ is the axial resolution, $\Delta\lambda$ is the FWHM source bandwidth, $\lambda_o$ is a central wavelength, and $\Delta k = (2\pi\cdot\Delta\lambda)/(N\cdot\lambda_0^2)$ is the wavenumber width at a single pixel in the line array camera for FD-OCT. The coefficient σ depends on the spectral shape, with a minimum of σ = 2ln2/π = 0.44 for a Gaussian spectrum and larger values otherwise, up to σ = 0.62 for a rectangular (flat-topped) spectrum.

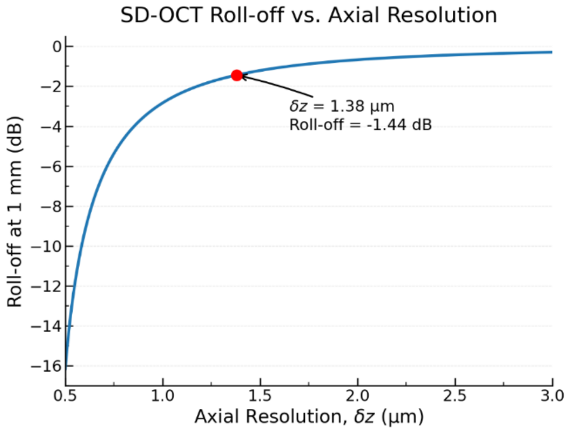


Figure 2. Theoretically limited roll-off as a function of the axial resolution of the FD-OCT system with a Gaussian spectrum, following Eq. 3. The red dot marks the parameters of the FD-OCT system used in this manuscript.

As shown in Fig. 2, the pixel sampling roll-off worsens steeply as the axial resolution improves. Finer axial resolution requires a broader wavenumber band per pixel, which sharpens the sinc envelope and accelerates depth decay. Even in the absence of optical aberrations, therefore, the pursuit of higher axial resolution in FD-OCT entails a progressively more severe roll-off and a correspondingly shorter usable imaging depth. This intrinsic trade-off represents a fundamental limitation of spectrometer-based detection in FD-OCT, independent of spectrometer quality, and is one that swept-source detection is not subject to.
